# Simulated-to-real Benchmarking of Acquisition Methods in Metabolomics

**DOI:** 10.1101/2023.01.12.523759

**Authors:** Joe Wandy, Ross McBride, Simon Rogers, Nikolaos Terzis, Stefan Weidt, Justin J.J. van der Hooft, Kevin Bryson, Rónán Daly, Vinny Davies

## Abstract

Data-Dependent and Data-Independent Acquisition modes (DDA and DIA, respectively) are both widely used to acquire MS2 spectra in untargeted liquid chromatography tandem mass spectrometry (LC-MS/MS) metabolomics analyses. Despite their wide use, little work has been attempted to systematically compare their performance due to the difficulty and cost of performing comparisons with real experimental data due to the lack of ground truth and the costs involved in running large number of acquisitions. Here, we present a systematic *in-silico* comparison of these two acquisition methods. To do so, we extended our Virtual Metabolomics Mass Spectrometer (ViMMS) framework with a DIA module. Our results show that the performance of these methods varies with the average number of co-eluting ions as the most important factor. At low numbers, DIA outperforms DDA, but at higher numbers, DDA has an advantage as DIA can no longer deal with the large amount of overlapping ion chromatograms. Results from simulation were further validated on an actual mass spectrometer, demonstrating that using ViMMS we can draw conclusions from simulation that translate well into the real world. Embedding this work within ViMMS also allows for researchers to easily simulate DDA and DIA LC-MS/MS runs, validate them on actual instrument, and potentially prototype novel methods that best combine the characteristics of both approaches. We believe that this work provides a useful guide to the choice of DIA or DDA for scientists involved in metabolomics data acquisition.

## 1 Introduction

Liquid chromatography tandem mass spectrometry (LC-MS/MS) is one of the dominant analytical platforms for untargeted metabolomics. LC-MS/MS acquisition strategies can be categorised as either Data-Dependent Acquisition (DDA) or Data-Independent Acquisition (DIA). In the former, MS2 scans are scheduled to target particular ions observed in full scan (MS1) survey scans. After each MS1 survey scan, a small number of ions will be prioritised (normally based upon their intensity) and fragmented in a series of MS/MS (MS2) scans. This will be followed by another MS1 survey scan from which the next batch of fragmentation events will be decided. In DIA, fragmentation is not based upon ions observed in survey scans but instead fixed m/z windows are isolated and fragmented, regardless of the ions present. The m/z windows can range from the whole m/z range (All-Ion Fragmentation, or AIF) or can be broken into a series of smaller windows that are iterated over in consecutive scans (e.g. Sequential Window Acquisition of all Theoretical Mass Spectra, or SWATH) [Sidoli et al., 2015]. DIA therefore removes the need to choose which ions to target at acquisition time at the cost of introducing an additional deconvolution step into the data analysis pipeline.

Various different DDA and DIA strategies have been introduced [Guan et al., 2020, Guo et al., 2021, Davies et al., 2021,Kaufmann and Walker, 2016] and although each new method is compared with other approaches, no clear consensus has emerged as to which overall strategy is best in which situation [Guo and Huan, 2020a,b, Fernández-Costa et al.,2020]. An advantage of DDA is that MS2 spectra are generated nearly ready to use, as each MS2 spectrum targets a particular ion, and we can be reasonably confident that the fragment ions observed do indeed come from the targeted ion. Sometimes multiple ions can end up in the same isolation window [Lawson et al., 2017], but this, in general, is not considered to be a major problem, as most of the times there is one dominant ion species giving rise to the mass fragments. Critics of DDA point to the lack of reproducibility (i.e., due to its stochastic nature, different peaks will be fragmented if the same sample is injected twice) and the low coverage – only a subset of the ions present in the sample are fragmented [Zhang et al., 2020]. As DIA does not prioritise based on the contents of MS1 survey scans, it is more reproducible (we know beforehand exactly the properties of any scan). It also, at least in theory, overcomes the coverage issues as it is able to assign MS2 fragments to any detected MS1 ion. However, because multiple ions will be fragmented in each MS2 scan, MS2 spectra must be deconvoluted before they can be used using software such as MS-DIAL [Tsugawa et al., 2015]. Deconvolution is not straightforward and it is widely recognised that the high coverage in DIA comes at the cost of lower quality spectra Bern et al. [2010].

Although there have recently been two studies [Guo and Huan, 2020a,b] that compare DDA with DIA for its use in metabolomics, in general comparisons between the two acquisition strategies are still lacking. This is mainly due to the lack of ground truth for real experimental matrices and the cost of running large numbers of injections. While validation on real injections is vital, simulation can also play an important role in answering such questions. For example, the number of ions that can be targeted in a DDA analysis is a complex function of scan times, chromatographic peak widths, and the number of peaks eluting at a particular time. Similarly, the number of spectra that can be accurately deconvoluted in a DIA analysis depends on the number of co-eluting ions in the same isolation window, and how correlated their chromatographic profiles are. In both cases, analysis on real injections is hampered by a lack of knowledge of the true make-up of that sample, or samples being overly simplistic if they consist of just a handful of known standards. Simulation can overcome these limitations by permitting complete control over the ground truth in terms of both the fragment spectra present, the number of chemical ions in the sample, and how and when they elute.

In our previous work we introduced ViMMS [Wandy et al., 2019, 2022], a virtual metabolomics mass spectrometry simulator framework, and demonstrated how it could be used to develop new, and improve existing DDA strategies. One of the main advantages of using ViMMS to develop new strategies lies in its ability to develop methods without the overhead costs of using a real mass spectrometer (MS). Within the ViMMS framework, it is possible to prototype methods and optimise them *in-silico* before transferring the developed method for validation to an actual instrument. New DDA methods, such as WeightedDEW and SmartROI, have been developed on top of ViMMS and shown to outperform Top-N in terms of the number of peaks that were fragmented in both simulated and real experiments [Davies et al., 2021]. More recent work has used ViMMS to develop improved methods for multi-sample and -injection DDA strategies [McBride et al., 2022].

In the current work we utilised ViMMS as a method for accurately benchmarking DIA and DDA through simulations. We firstly introduce two DIA controllers (SWATH and AIF) into the ViMMS framework, before conducting extensive simulated experiments to evaluate the comparative performance of DIA and DDA (Top-N) across a range of different simulated conditions, effectively creating a “digital twin” of the real situation. The DIA methods prototyped on the simulator were transferred with ease to run on an actual MS instrument with no code changes in the implementation – evidencing the capability of ViMMS to develop DIA methods in a simulated-to-real setting. As such, we used ViMMS to validate the *in-silico* experiments using complex beer samples in real LC-MS/MS experiments. The real experimental results were first benchmarked using an online reference spectral library (GNPS/NIST14) to assess spectral matches to a database of known molecules. This approach follows that of Guo and Huan [2020a], but importantly compares SWATH as well as AIF. Improving upon Guo and Huan [2020a], we also more systematically compare the results for DDA (Top-N) and DIA (SWATH and AIF) by evaluating our results against a database created on the specific sample using a recently developed, but computationally expensive multiple injection data acquisition method [McBride et al., 2022]. This multi-injection dataset was constructed specifically to evaluate the maximum spectral coverage from a realistic experimental setting, complementing the standard approach of evaluating against a database of known molecules.

Our results found that over a wide range of experimental conditions, DIA is generally more effective at fragmenting more features, both in simulations and reality. However, DDA outperforms DIA in terms of the number of chemical ions for which high-quality spectra are recovered. Based on simulated and real instrumental results, we were able to provide a clear, actionable guideline on when a particular acquisition method (whether DDA or DIA) should be used. In summary, the contributions of this paper are as follows:

1. We have introduced two new DIA controllers (SWATH, AIF) into the ViMMS framework.
2. We have conducted extensive simulated experiments to evaluate the performance of DDA (Top-N) vs DIA with a known *in-silico* ground truth.
3. We have validated the simulated results through benchmarking on the actual instrument using two reference datasets: the GNPS/NIST14, and our own Multi-Injection libraries.

## 2 Materials & Methods

### 2.1 DDA and DIA Data Acquisition

To validate performance on a real instrument, we performed DDA and DIA acquisition using six beer samples. Each beer sample was acquired once using the Fullscan, Top-N, AIF and SWATH controllers in ViMMS when connected to an actual mass spectrometer (more details in Section 2.4). To create the Multi-Injection reference library (described in Section 2.5), each beer sample was further injected ten times repeatedly for acquisition using the Intensity Non-overlap method in ViMMS [McBride et al., 2022].

For sample extraction, chloroform and methanol were added to beer samples (detailed names in Supplementary Section S1) in a 1:1:3 ratio and mixed with a vortex mixer. The mixture was centrifuged to remove protein and other precipitates, and the supernatant was stored at −80° C. Chromatographic separation was performed with a Thermo Scientific UltiMate 3000 RSLC liquid chromatography system and a SeQuant ZIC-pHILIC column. The gradient elution used (A) 20 mM ammonium carbonate and (B) acetonitrile. 10 *μ*L of each sample was injected with an initial 80% acetonitrile concentration (B), maintaining a linear gradient from 80% to 20% (B) over 15 min, and finally a wash of 5% (B) for 2 min, before re-equilibration at 80% (B) for 9 min. The flow rate was 300 μL/min and the column oven temperature was 40° C.

A Thermo Orbitrap Fusion tribrid-series mass spectrometer was used to generate mass spectra data, controlled through Thermo Instrument Application Programming Interface (IAPI) managed by ViMMS (more details in Section 2.4). Full scan spectra were acquired in positive mode at a resolution of 120,000 and a mass range of 70-1000 m/z. Fragmentation spectra for both DDA and DIA were acquired using the Orbitrap mass analyser at resolution 7,500. In DDA mode, precursor ions were isolated using 0.7 m/z width and fragmented with fixed HCD collision energy of 25%. The AGC was set as 200,000 for MS1 scans and 30,000 for MS2 scans. *N* was set to 10 Top-N. The dynamic exclusion window (DEW) was set to 15s to prevent repeated fragmentation of the same ion. A minimum intensity threshold of 5000 was also used before a precursor ion can be selected for MS2 fragmentation. For DIA (AIF), an MS1 source CID energy of 25% was used. For DIA (SWATH), a window of 100 m/z was used with no overlap between the windows.

### 2.2 Developing DIA Methods Using ViMMS

In previous work [Davies et al., 2021], ViMMS was used to develop DDA methods, but the capability of the framework is not limited to that. Here we introduced two new methods, SWATH and AIF, on top of the framework, demonstrating that DIA methods can also be developed on top of ViMMS. Rather than prioritising ions for fragmentation based on their abundance as is commonly done for DDA, DIA methods operate by fragmenting all precursor ions within a large m/z window. In AIF, all precursors in the entire m/z range are fragmented, whereas in SWATH, a series of smaller and potentially overlapping windows are used to fragment ions in the window. Both AIF and SWATH were implemented as controllers in ViMMS, allowing their performance to be benchmarked in the simulator and validated on an actual instrument easily, as previously done with DDA methods [Davies et al., 2021].

Figure 1 shows a schematic of the DIA method implementations in ViMMS illustrating how the new DIA methods were introduced. The ViMMS framework can be divided into two parts: a ‘Simulated Environment’ where simulated scans are generated by querying synthetic chemicals, and a ‘Real Environment’ where actual scans are generated through measurements using an LC-MS instrument. In the Simulated Environment (Figure 1A), the Virtual MS is seeded with synthetic molecules that are created by either sampling chemical databases or extracted from existing experimental mzML files. Once generated, molecules can be used to produce scans during virtual mass spectrometry. Scans are generated based on which molecules elute at a particular retention time, and generated chemicals are dispatched to the appropriate controllers. The new DIA methods are implemented as the SWATH and AIF controller classes in Python (solid purple box in Figure 1A), extending from the base Controller class in ViMMS (blue box in Figure 1A). Controller classes follow a predefined Python interface to receive scans and schedule the next MS1 and MS2 scans. Different experiments can be performed in the Simulated Environment, allowing for different data characteristics to be explored. Once tested and optimised, the developed DIA controllers can be transferred to run on an actual mass spectrometer instrument with no change to their implementations. This is accomplished by swapping the Simulated Environment to a Real Environment (Figure 1B), where an IAPI MS is used in place of the Virtual MS. The IAPI MS class has the same interface as the Virtual MS to ensure code compatibility, however the IAPI MS relies upon the Instrument Application Programming Interface (IAPI) [“Thermo Fisher Scientific”, 2022] to communicate with an actual Thermo Orbitrap Fusion instrument. All developed controller implementations, whether DDA or DIA that were initially tested against the Simulated Environment in ViMMS, can run without any change in the Real Environment (for the new DIA methods, this is the dashed purple box in Figure 1B). In this manner, real experimental data can be seamlessly acquired using the AIF and SWATH controllers initially developed in the simulator.

**Figure 1:**
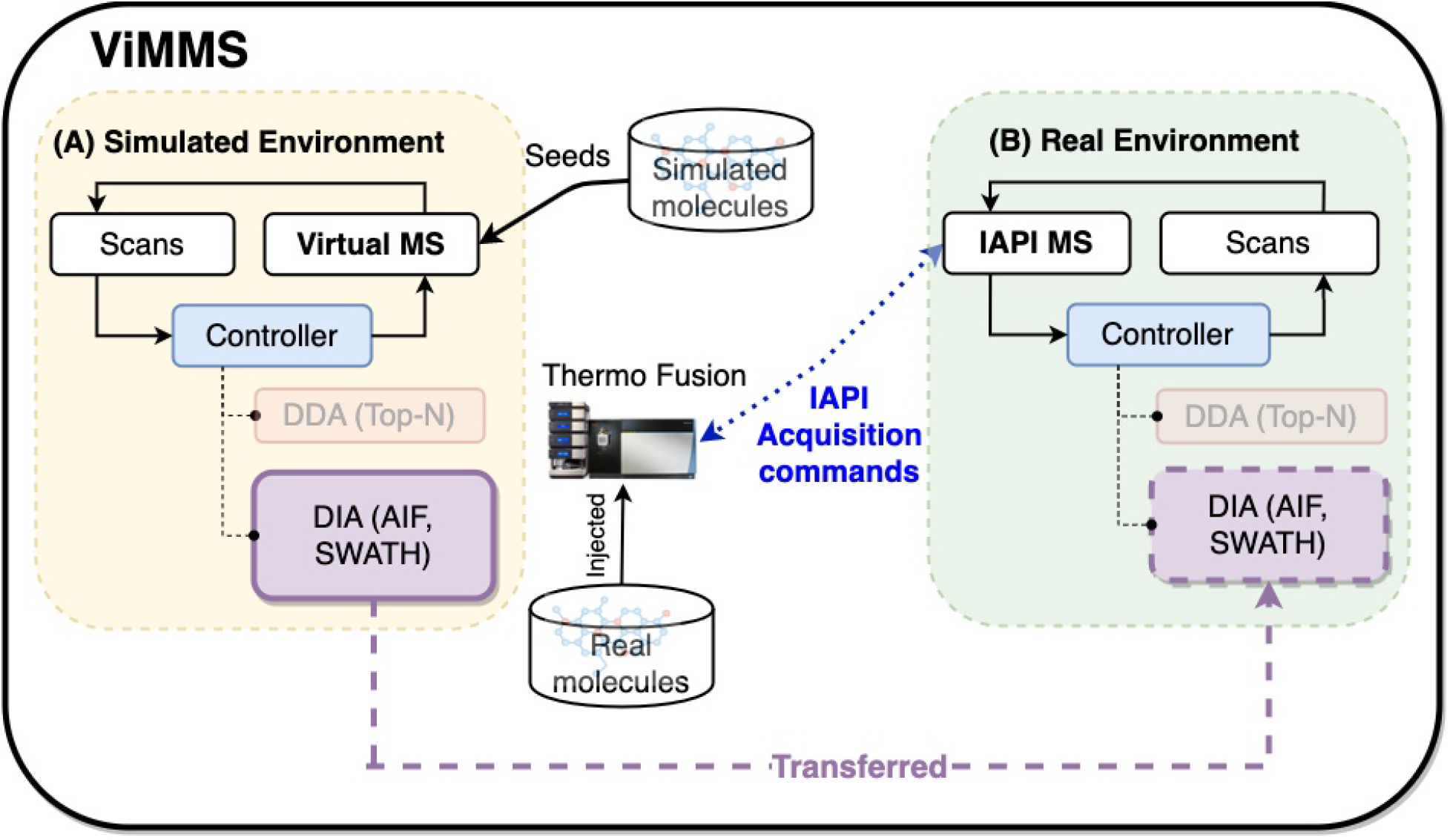
The overall schematic of the ViMMS framework. **(A)** The Simulated Environment in ViMMS allows for new acquisition methods to be developed against a Virtual MS that takes simulated molecules as input. The new DIA methods, e.g. SWATH and AIF (solid purple box), as well as existing DDA methods (faded orange box), are implemented as controllers in Python, and initially tested here in the Simulated Environment. **(B)** Acquisition methods can be run for method validation on the Real Environment in ViMMS, connected to a Thermo Fusion instrument via IAPI, to acquire real experimental scans. The underlying Python controller codes for the new DIA methods remain unchanged when transferred from the Simulated to the Real Environment (dashed purple box).

### 2.3 Simulating DDA and DIA Methods

#### 2.3.1 Generation of Simulated Data

Simulations allow us to flexibly define different scenarios to validate hypotheses without costly instrument time. To compare the two types of acquisition methods, we use the Simulated Environment in ViMMS (Figure 1A) to generate simulated data with an increasing number of co-eluting chemicals present to test the limit of deconvolution empirically. While these chemicals are purely synthetic, they allow us to know the ground truth and evaluate the results of the different methods more accurately without being reliant on inconsistent database matching.

In the simulated study, each chemical is generated by first choosing a formula randomly from the HMDB database, ensuring that its observed monoisotopic mass is between 100-1000 Da. Next the chemical is assigned a uniformly-sampled retention time value between 0 - 400s. The choice of uniform distribution here is motivated by the narrow retention time range used. When the number of chemicals are high (e.g. 5000), this results in dense regions of co-eluting ions throughout the entire simulated injection, which could challenge spectral deconvolution and potentially demonstrate the limit of such approaches.

Finally for each chemical, a chromatogram is generated. The maximum intensity value of the apex of the chromatogram was sampled from a uniform distribution between 1E4 - 1E7. A Gaussian chromatographic peak shape with a mean centered at the chemical’s RT value, and a standard deviation of 5s is assumed. This chromatographic peak shape assumption matches that of MS-DIAL, thus making the task of peak picking easier. Each chemical’s observed MS2 spectra are also generated randomly by sampling fragment peaks’ m/z uniformly between 70 m/z to the exact mass of the formula (assuming a positive charge of +1). The intensity of a fragment peak is also set to be between the specified minimum (0.1) and maximum (0.8) proportions of the chemical’s apex intensity. The number of fragment peaks in an MS2 scan is generated by sampling from a Poisson distribution with the mean 10. This Poisson mean was chosen to produce sufficient expected numbers of fragment peaks per MS2 scan for spectral matching later.

Simulated samples containing the specified number of chemicals are generated in a case-vs-control setup, where each sample set consists of 5 case and 5 control samples, and each sample contains observation from the specified number of chemicals. To reduce the influence of peak picking and alignment during data processing of DIA data, chemicals were generated such that the same chemical has identical m/z and RT values across samples, although their intensity values change across the case and control groups. While simple, this setup represents a realistic case that captures the essence of many real biological mass spectrometry-based experiments. Varying numbers of chemicals are generated during simulation, ranging from 10, 20, 50, 100, 200, 500, 1000, 2000, 5000. The entire experiment is repeated 5 times, resulting in multiple experimental replicates. DDA (Top-N) and DIA (AIF, SWATH) controllers were run for every sample that was simulated, resulting in an mzML file for each acquisition run. Parameters of these controllers were set to be the same as that used for real data acquisition (detailed in Section 2.1.

#### 2.3.2 Processing of Simulated Data

Once simulated data has been generated in Section 2.3.1, chemicals need to be mapped to their respective fragmentation spectra. For DDA, the association between observed fragmentation scans to chemicals is known unambiguously as their mapping can be read from the simulation state directly. For DIA, spectral deconvolution needs to be performed to assign the deconvoluted fragment peaks to chemicals. We chose MS-DIAL [Tsugawa et al., 2015] for peak picking and spectral deconvolution on the resulting mzML files produced by simulating DIA methods (see Supplementary Section S2 for the MS-DIAL parameter settings). MS-DIAL was chosen as it was widely used by the community for processing DIA data.

From MS-DIAL output, we can extract for each sample set a list of features that were detected and aligned across the DIA mzML files (5 cases and 5 controls). Each feature is potentially associated with a fragmentation spectra, which has been deconvoluted by MS-DIAL. To assign fragmentation spectra, we matched simulated chemicals to features using the m/z tolerance of 5 ppm and RT tolerance of 10 seconds, therefore linking chemicals to their deconvoluted spectra. If there are multiple possible candidates during matching, the feature closest in m/z value to the chemical’s monoisotopic peak will be chosen. Finally for each simulated chemical, its true fragmentation spectra are known. This can be used to construct a library of true reference spectra for matching. Observed (and deconvoluted, in the case of DIA) spectra are matched to the true spectral library using cosine similarity. For matching, a bin width of 0.05 Da is used and a minimum of at least 3 matching peaks is required.

### 2.4 Validation on a Real Instrument

#### 2.4.1 Generation of Real Data

The Real Environment in ViMMS was used to validate that the simulated results translate to real experiments (Figure 1B). This environment can be connected to a Thermo Orbitrap Fusion tribrid-series mass spectrometer, allowing us to perform data acquisition on a series of beer samples using the fullscan, Top-N, and the newly introduced SWATH and AIF controllers. The result from data acquisition is a series of mzML files, one for each beer sample and controller used. For more details on real data acquisition, refer to Section 2.1.

#### 2.4.2 Processing of Real Data

Peak picking and alignment was performed using MS-DIAL on the real experimentally-derived fullscan mzML files. Unlike in simulations, here the exact chemical composition of the sample is unknown. As such, fullscan features from peak picking on the fullscan data are used as a proxy for chemicals. The fullscan data, which contains the most MS1 information and therefore the best chromatographic peak shapes, was chosen for feature extraction using MS-DIAL. MS-DIAL parameters were chosen by hand to give a reasonable number of peaks comparable to what we have seen from past experiments using this kind of sample on the same instrument (details in Supplementary Section S3). The result from this is a list of fullscan features detected and aligned from the fullscan beer mzML files.

Next DDA fragmentation spectra were to be assigned to fullscan features, with the following procedure employed to bypass the need to do further peak picking on the DDA mzML files (which can be problematic due to the lower number of MS1 scans in fragmentation files). *pymzML* [Bald et al., 2012] was used to load MS2 scans from each DDA mzML file. For each MS2 scan, we used its precursor m/z, isolation window and RT values to assign the scan (and therefore fragmentation spectra) to its corresponding fullscan feature. This assignment was done based on whether the isolation window and RT values of a chemical overlap with the feature’s bounding box from peak picking. If a fullscan feature had multiple MS2 scans associated with it, the scan that was fragmented at the highest intensity within the feature’s bounding box was chosen.

Finally, just like in simulation, MS-DIAL was used to perform peak picking and spectral deconvolution on the DIA mzML files. Peak picking was performed using the same parameters as the fullscan data, and an additional deconvolution step was done using MS-DIAL (detailed parameters in Supplementary Section S3). From MS-DIAL output we extracted a list of fragmentation features detected in the DIA mzML files and their corresponding MS-DIAL deconvoluted spectra. Fullscan features (without fragmentation information) were matched to the DIA fragmentation features (with deconvolved spectral information) using an m/z tolerance of 5 ppm and RT tolerance of 10 seconds. If there were several possible candidates during matching, the one closest in m/z value was chosen. The results of this procedure from both the DDA and DIA data is the assignment of a fullscan feature to fragmentation spectra. Such spectra can be used for matching fullscan features to the reference libraries during benchmarking experiments.

### 2.5 Matching Experimental Spectra to Reference Libraries

To assess mass spectral quality, we match the fragmentation spectra from DDA and DIA methods to two reference spectra libraries. These reference libraries can be used to match against the observed and deconvoluted spectra from DDA and DIA methods. For matching, a bin width of 0.05 Da is used and a minimum of at least 3 matching peaks is required.

The first reference library is the ‘GNPS Matches to NIST14’ dataset obtained from the GNPS library [Wang et al.,2016]. This dataset contains 5763 high confidence matches to NIST14 MS/MS library spectra. Filtering by polarity is performed to select spectra in positive mode only that can be used as reference to assess spectral annotation quality. For experiments, we call this the *GNPS/NIST14* library. Its purpose is to assess how many potentially unknown metabolites could be annotated in an untargeted metabolomics experiment using the different acquisition methods.

Additionally, we also introduced our own multiple-injection reference library for spectral matching, which we call the *Multi-Injection* Library. Each beer sample was injected ten times and data acquisition is performed by taking advantage of replicate information using the Intensity Non-overlap method available in ViMMS (detailed in Supplementary Section S4). Intensity Non-overlap is an iterative DDA-based method that detects regions-of-interests (ROIs) in real-time and avoids re-fragmenting the same ROI multiple times across successive injections. An ROI is scored for fragmentation based on its overlapping area weighted by intensity, with higher scoring ROIs selected more often (for more details, refer to Supplementary Section S4 and McBride et al. [2022]). In the presence of multiple injections, Intensity Non-overlap has been shown to outperform Top-N by a large margin as it is able to target more unique features across injections, while fragmenting each feature closer to its apex.

To convert the acquired Multi-Injection mzML files into spectral library, for each beer sample, we perform peak picking using MS-DIAL on its corresponding fullscan mzML file, generating a list of fullscan feature for that beer sample. Fragmentation spectra, acquired via exhaustive fragmentation of each sample using Intensity Non-overlap, were extracted from mzML files and matched to the detected fullscan features following the DDA procedure outlined in Section 2.4.2. The purpose of introducing this Multi-Injection reference library is to assess the coverage of the benchmarked DDA (Top-N) and DIA (AIF, SWATH) methods when only a single replicate is available (a common occurrence) and increasing the number of replicates is not possible due to cost or other constraints.

## 3 Results

### 3.1 Simulated Results

Acquired fragmentation spectra could help deduce the chemical identities of measured compounds. From the proposed simulated experiment in Section 2.3, for each acquisition method the number of chemicals that could be annotated based on spectral matching is obtained. The number of unique chemical annotations in the dataset is counted at varying thresholds for cosine similarity score of at least 20%, 40%, 60% and 80% (Figure 2). The results across 5 replicates show that for small numbers of chemicals, all benchmarked methods, whether DDA or DIA, have similar annotation performance. At similarity threshold 60% and with only 100 chemicals, AIF annotated a mean of 72.6% of chemicals, SWATH 86.0% and Top-N 84.4%. As the number of chemicals increases, the gap between the benchmarked methods widen leading to lower annotation rates from DIA. When the number of chemicals increased to 200, at 60% similarity threshold, AIF managed to annotate 54.4% chemicals, SWATH 80.7% and Top-N 82.4%. For 500 chemicals, AIF annotates 28.8% of chemicals, SWATH 59.0% and Top-N 79.3%. With a greater number of chemicals, Top-N outperforms the two DIA methods across all thresholds. At the highest number of chemicals (5000), the annotations obtained from SWATH and AIF are nearly 0% while Top-N managed a mean of 31.5% across replicates. It can be observed that increasing the cosine similarity threshold from 20% to 80% lowers the results of all methods, but the overall trend remains.

**Figure 2:**
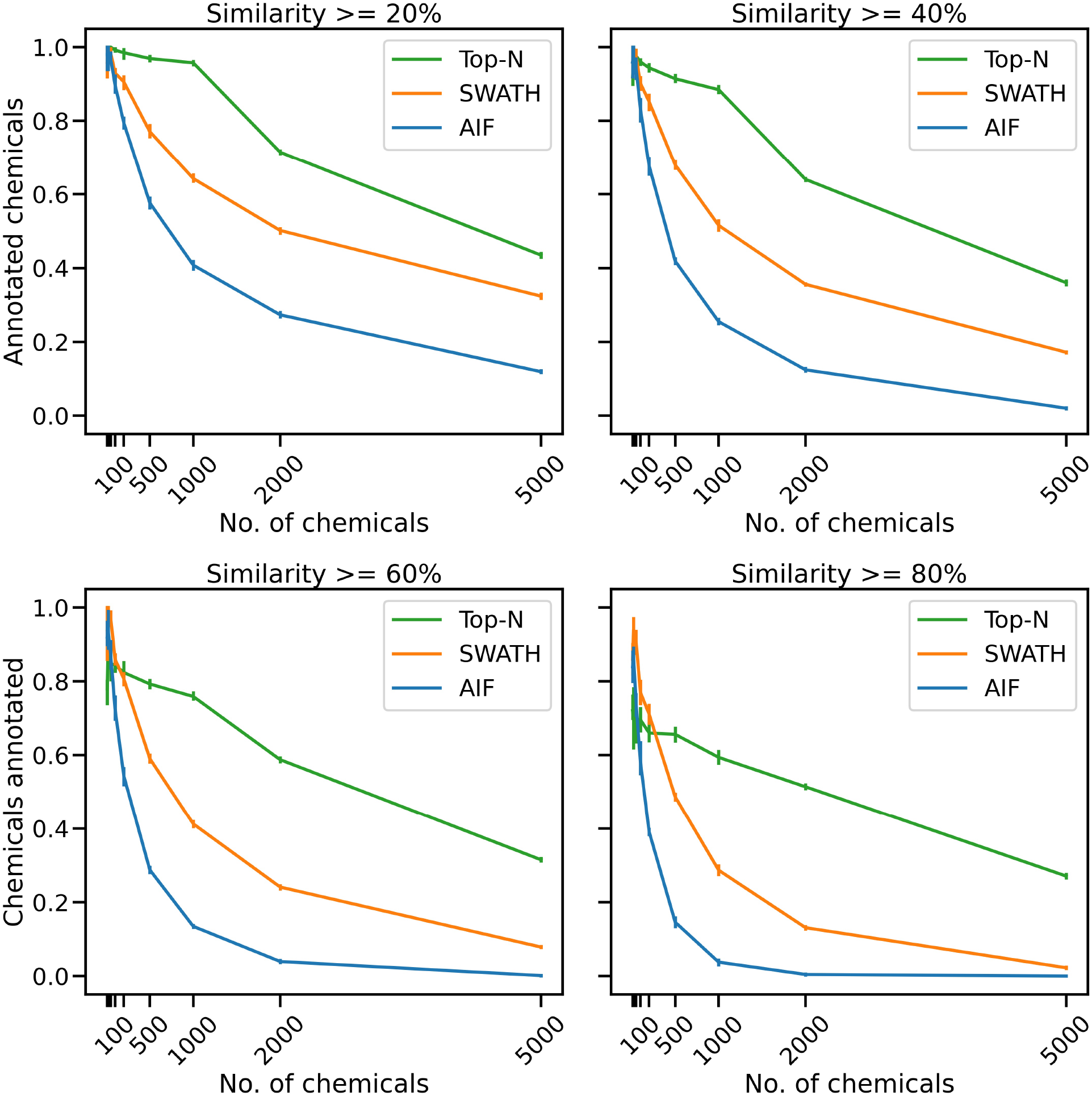
The mean proportion of unique chemical annotations at varying numbers of chemicals and similarity thresholds across 5 replicates. The error bar shows the 95% confidence interval.

To explain the annotation results in Figure 2, here we consider for one replicate the distribution of spectral similarity scores when matching DDA and DIA spectra to the known ground truth of true chemical fragmentation spectra. The results are shown in Figure 3 for the entire range of chemicals tested. From Figure 3, it can be seen that Top-N generally performs best in returning high similarity scores, followed by SWATH then AIF. For Top-N, the cosine similarities of matches generally remain high even with an increasing number of chemicals. For the two DIA methods, their cosine similarities gradually decrease with more co-eluting chemicals. This could be explained by the fact that as the elution profile gets more crowded, more precursor ions are isolated and fragmented in the same window, making deconvolution harder. SWATH performs marginally better than AIF with increasing chemicals. This makes sense as the window used for isolating multiple peaks in SWATH is smaller than AIF where all ions in the entire scan range are used, making the deconvolution problem slightly easier for SWATH.

**Figure 3:**
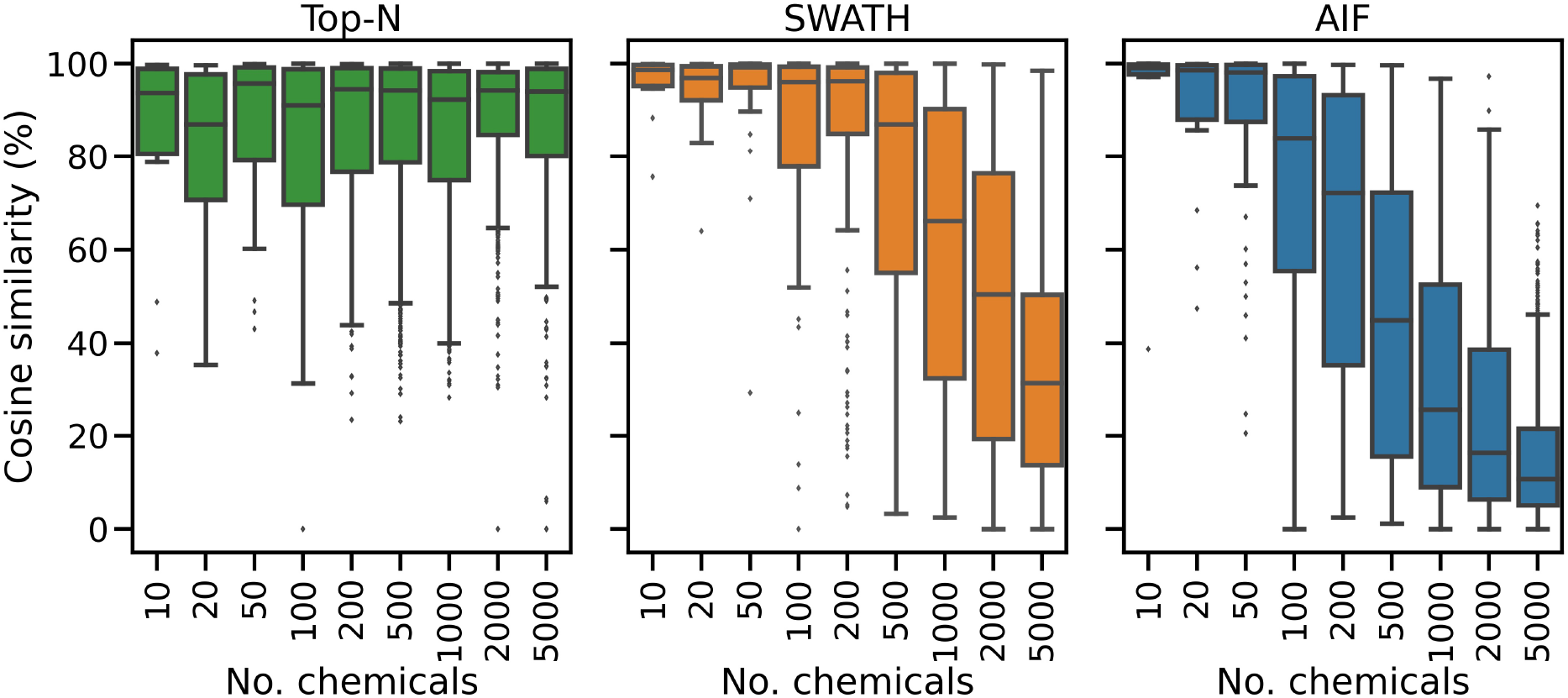
The distribution of cosine similarity scores when matching observed spectra to the true reference spectra for varying numbers of chemicals.

Inspecting the hardest case of 5000 chemicals in Figure 4, it can be observed that AIF returns most of its matches at low cosine similarity achieving a median score of 10.7%, Top-N returns most matches at high cosine similarity with a median of 94.1%. SWATH outperforms AIF (median similarity 31.3%) but not as well as Top-N. To explain the decrease in similarity scores, we inspect the pairwise cosine similarity of all observed/deconvolved spectra for each acquisition method (Figure 5). The results show that the pairwise similarity of the ground truth and Top-N spectra is nearly 0 for nearly the entire range of chemicals. However, most likely due to the increased difficulty in deconvolution, pairwise spectral similarities in the SWATH and AIF results are higher with increasing chemicals – resulting in a decrease in identification hits, and fewer matches at high cosine similarity as the experimental mass spectra become more similar to each other and less similar to the true mass spectra of the chemicals.

**Figure 4:**
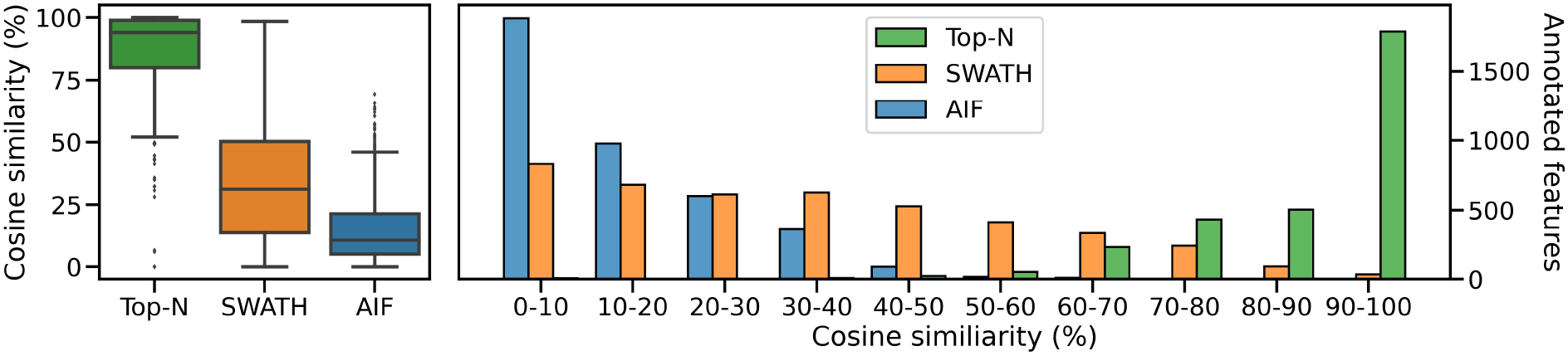
The distribution of cosine similarity scores as a boxplot (left) and a histogram (right) when matching observed spectra to the true reference spectra for 5000 chemicals.

**Figure 5:**
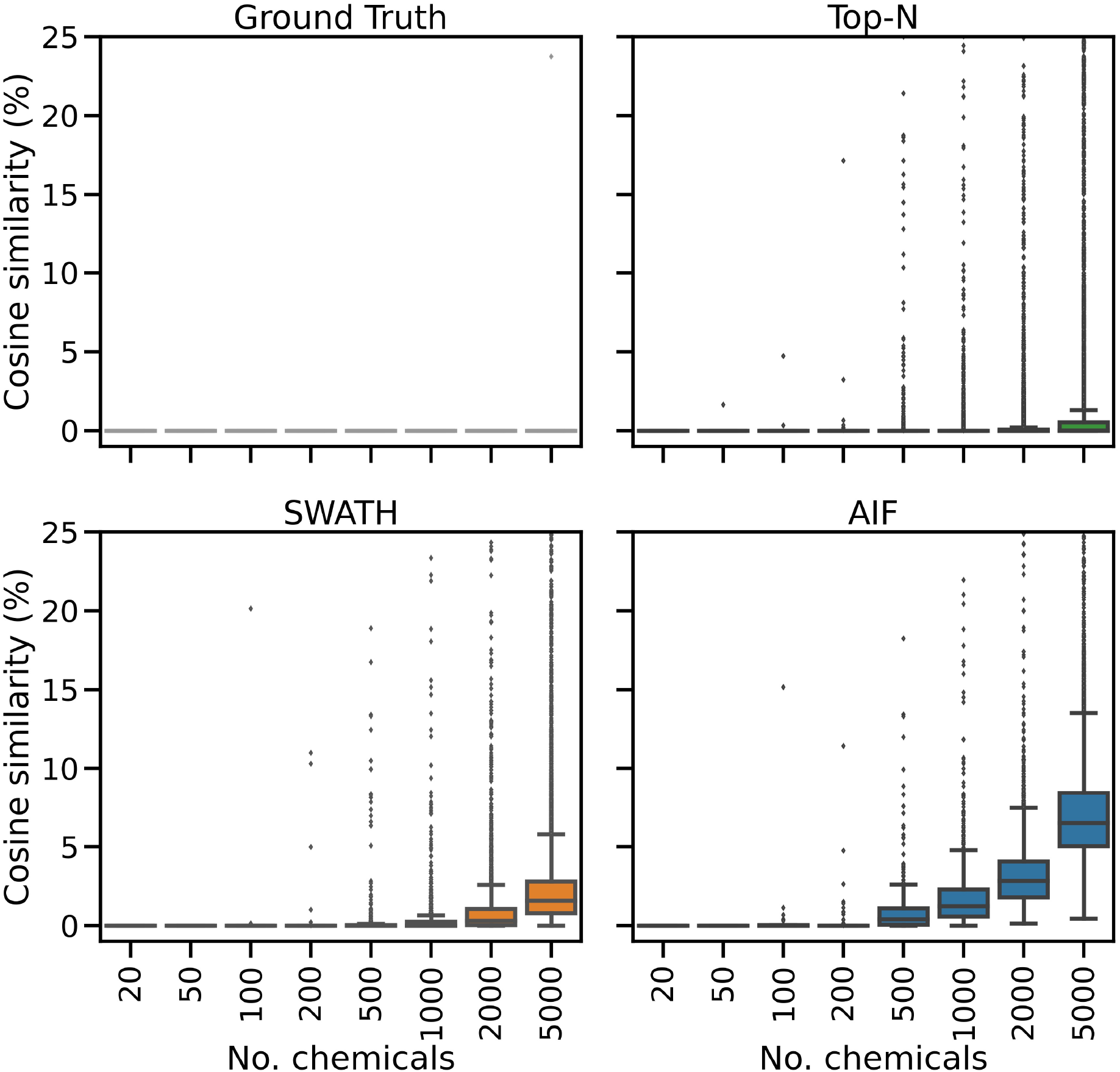
The distribution of pairwise cosine similarity scores of the ground truth (true chemical spectra), and fragmentation spectra from Top-N, SWATH and AIF. The plot y-axes are truncated at 25% similarity.

### 3.2 Real Experimental Results

#### 3.2.1 Mapping Features to Fragmentation Spectra

Simulated results were further validated on real instruments by running the benchmarked DDA and DIA methods on actual beer samples. Following the procedure in Section 2.4.2 to map fullscan features to fragmentation spectra, 6090 fullscan features initially were detected from the fullscan data after peak picking and alignment. After matching the fullscan features to fragmentation data, DIA methods produce the largest number of matched features, with SWATH at 3889 and AIF at 4381. In contrast, Top-N only has 2895 matched features. Figure 6 summarises the proportion of fullscan features that can be matched to the fragmentation spectra. Our results here agree with Guo and Huan [2020a] in how DIA (AIF) is able to fragment more features than DDA (Top-N). We note some slight differences in our methodology to [Guo and Huan, 2020a]. In this work, we perform peak picking to extract features on the fullscan data, which has more reliable MS1 signals. Fragmentation mzML files are used only to map fragmentation scans to the detected fullscan features. whereas in [Guo and Huan, 2020a] the peak picking is performed directly on the fragmentation mzML files, potentially leading to poorer extracted chromatographic peak shapes due to sparser MS1 data points in the mzML files. Despite these differences in the experimental set up, the conclusions from both our study and [Guo and Huan, 2020a] agree that AIF outperforms Top-N in the number of fragmented features. Additionally SWATH was not included in [Guo and Huan, 2020a] but it was hypothesised to perform in the middle of Top-N and AIF, as the windowing approach used in SWATH lies between the two extremes of fragmenting everything (AIF) and fragmenting only a few selected precursor ions (Top-N). Our results here confirmed that SWATH achieves a higher coverage than Top-N but lower than AIF.

**Figure 6:**
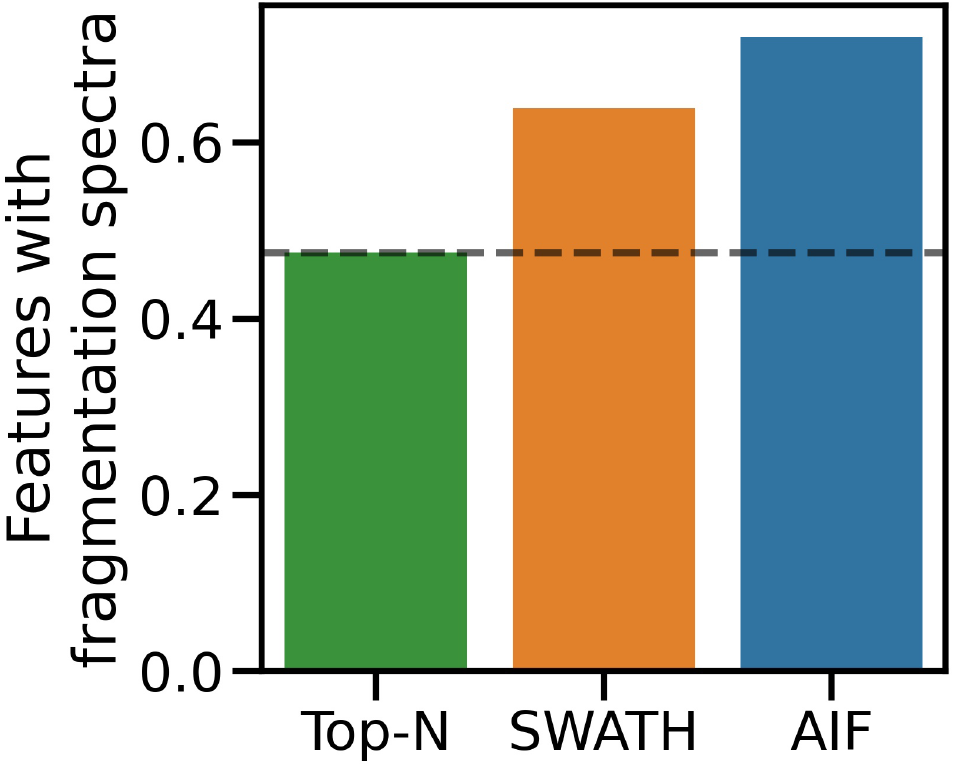
Proportion of matched features to the total number of detected features from the fullscan data.

#### 3.2.2 Pairwise Spectral Similarity

By following the methodology of Section 2.5, two reference mass spectral libraries were constructed, one based on the GNPS spectra matched to NIST-14 at high reliability (the GNPS/NIST14 library), and another based on an exhaustive multiple-injection approach we constructed ourselves (the Multi-Injection library). The purpose of the GNPS/NIST14 library is to assess how many unknown molecules can be identified from spectral matching for each acquisition method, whereas the purpose of the Multi-Injection library is to determine the extent of coverage with respect to an exhaustive and expensive multiple-injection method. After filtering by polarity, we obtain 5274 positive-mode spectra from the GNPS/NIST14 library, whereas for the Multi-Injection library, in total 4987 features were available for matching in this library.

To assess spectral quality, we first compute the pairwise similarity of spectra in the same dataset, and also within each of the two reference libraries. For each feature, we find the matches to fragmentation spectra in the same dataset by computing the cosine similarity (ms2_tol = 0.05 Da, min match peaks = 3), with the results shown in Figure 7. The Multi-Injection reference library has a median of nearly 0% for pairwise similarities, demonstrating that the acquired reference spectra using the Intensity Non-overlap method are sufficiently different from each other. The GNPS reference library has a higher pairwise similarity of 9.5%, suggesting that some of the curated molecules share similar structures and therefore similar mass fragmentation spectra. This is a reasonable assumption as shared structures is the core underlying assumption for models to discover chemical substructures, such as MS2LDA [van Der Hooft et al., 2016] that has been applied to reference libraries like MassBank and GNPS.

**Figure 7:**
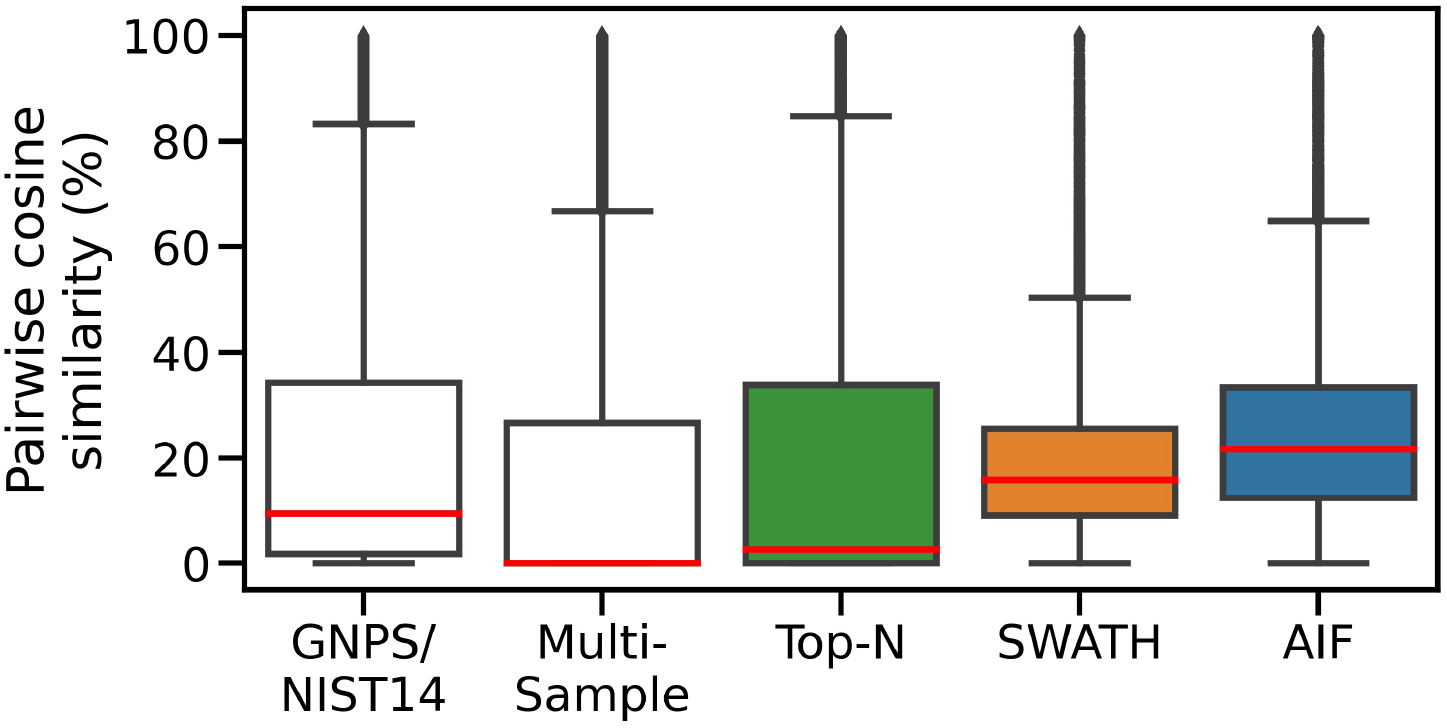
Pairwise similarities of spectra in each dataset. DIA methods (AIF, SWATH) produce deconvoluted spectra that are more similar to each other.

The Top-N dataset also has the same low median pairwise similarity (2.7%) in its member spectra. This is expected given the nature of DDA. Assuming that deconvolution works well for DIA data and the resulting spectra are dissimilar to each other in a manner similar to the Top-N results, we expect to observe lower pairwise similarities from the DIA datasets too. However the higher median pairwise similarity scores for SWATH (16%) and AIF (22%) suggest that deconvolved spectra from SWATH and AIF tend to be more similar to each other. This could be due to the difficulty in deconvolving spectra. The results here generally agree with simulated results in Section 3.1, where DIA methods are also shown to exhibit higher pairwise similarity in simulation.

#### 3.2.3 Spectral Matching Results

We compute the cosine similarity of features to the GNPS/NIST14 reference spectra. Similar to before, features were matched to reference spectra based on their cosine similarity (ms2_tol = 0.05 Da, min match peaks = 3). if there are multiple matches for each feature, the one with the best score (highest) is kept. Figure 8A shows the score distributions for the three methods. From Figure 8A (left panel), it can be observed that Top-N obtains the highest median cosine similarity (26.5%) followed by SWATH (19.5%) and finally AIF (15.2%). Consequently this results in Top-N (DDA) obtaining the most high-scoring matches at ≥ 60% similarity, followed by SWATH and AIF last (Figure 8A). The results here are consistent with the simulated results in Section 3.1 and also with Guo and Huan [2020a] but in that work, only Top-N and AIF were compared. Our results further confirmed the hypothesis in Guo and Huan [2020a] that SWATH should perform in the middle of Top-N and AIF when it comes to spectral quality. This is because deconvolution using SWATH data is easier than AIF due to the smaller window sizes.

**Figure 8:**
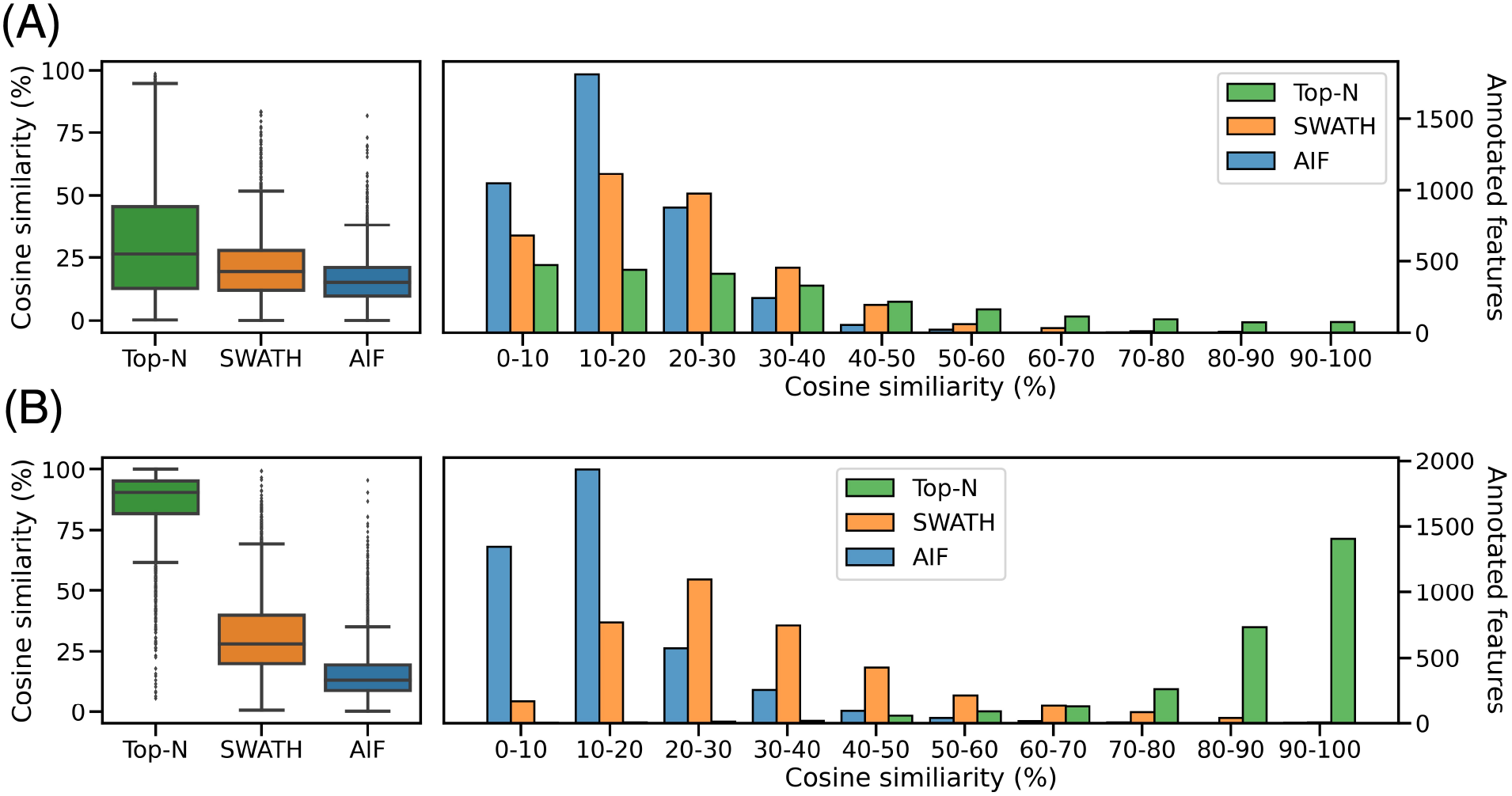
Distribution of cosine similarity of annotated features for Top-N and DIA methods for the **(A)** GNPS/NIST14 library and **(B)** Multi-Injection library.

Next we perform spectral matching to the Multi-Injection reference library. When matched against Top-N and DIA methods, Top-N performs best (90.5% median) followed by SWATH (28.1%) and AIF (13.3%) (Figure 8B). The results here are consistent with the GNPS results above. However we see that the median similarity of Top-N is much higher compared to the two DIA methods here. That is because the reference spectra contains more query spectra since it is based on the same sample – only fragmented exhaustively, and unlike GNPS/NIST14 there are fewer missing matches.

Finally we compare the overlap of annotated features at matching threshold ≥ 60% (Figures 9) for both reference libraries. Top-N (DDA) obtains the most hits, being able to annotate the most unique features (295 for GNPS/NIST14, and 2231 for Multi-Injection). In both cases, there is a large overlap between Top-N and DIA annotations, and most DIA annotations are also recovered by Top-N. Between AIF and SWATH, there is also a lot of overlap with SWATH being able to recover most of the annotations of AIF. The results here for AIF and Top-N generally agree with Guo and Huan [2020a], with SWATH a new addition in our results that perform in between those two.

**Figure 9:**
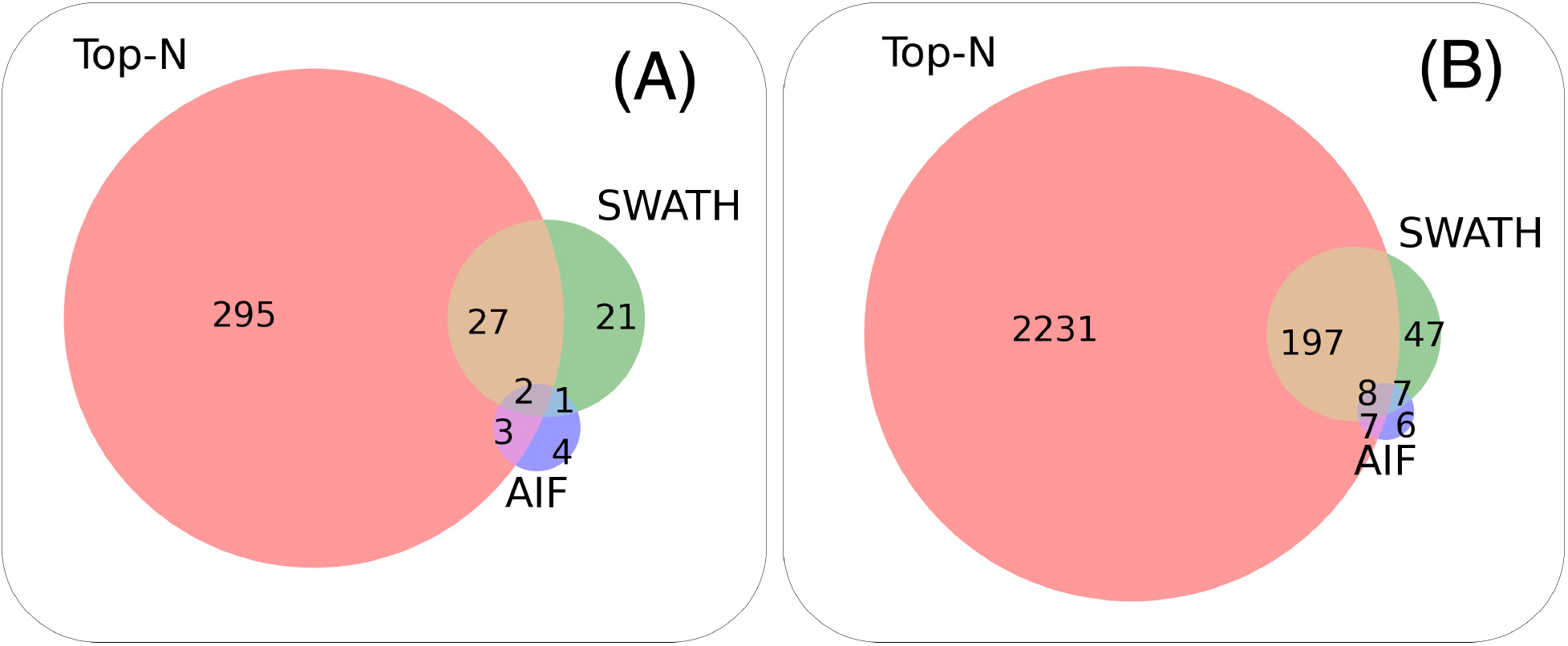
Venn diagram showing the overlap between Top-N, SWATH and AIF at matching threshold ≥ 60% for the **(A)**GNPS/NIST14 library, and **(B)**Multi-Injection library.

## 4 Discussion & Conclusion

In this study, we performed a comprehensive comparison of DDA vs DIA methods, first in the simulator, followed by validation using real experimental mass spectrometry data. We simulated experiments at various complexity levels to challenge DDA acquision methods and DIA deconvolution methods. Here, we observed that the quality of deconvolution using MS-DIAL is limited by the number of co-eluting chemicals. Identification performance is limited in DIA compared to DDA with a large number (more than 1000) of simulated chemicals or observed molecular features. We further benchmarked this scenario using a real untargeted metabolomics dataset acquired on the mass spectrometer generated from beer samples. Mass spectral matching of experimental mass fragmentation spectra from this real dataset on two sets of reference mass spectral libraries confirmed our simulated results. This validates our motivation that a simulator framework such as ViMMS can be used to benchmark the performance of both types of methods. Simulation helps to provide an environment that can be used to prototype and validate advanced deconvolution methods – without the need of costly instruments (i.e., a “digital twin”). This encourages the development of better deconvolution methods, since known ground truths were generated *in-silico*, thus making benchmarking easier. The ViMMS framework could also be used to simulate various scenarios that we have not covered in this study, for instance simulating different column properties and elution profile of chemicals, and assessing how that affects deconvolution.

In the context of LC-MS/MS analysis, a complex sample would contain a large number of different metabolites, while a simple sample would contain only a few metabolites. Determining the complexity of a sample is important for choosing the appropriate acquisition strategy for LC-MS/MS analysis. ViMMS can generate simulated samples of varying complexity, which can be used to assess whether DDA or DIA methods should be used in specific scenarios. By varying average number of co-eluting ions, we found that DIA being more effective at lower numbers and DDA having an advantage at higher numbers where DIA struggles to handle the large amount of overlapping ion chromatograms. From both simulated and real results, DDA was also found to generally perform better than DIA when it comes to matching unidentified features to spectra in both reference libraries (GNPS and Multi-Injection). DIA fragments more features than DDA but their quality for spectral matching is typically lower. Our results on this are not unique as a similar work in [Guo and Huan, 2020a] confirms our findings. Crucially, our study improves upon that prior work in several key aspects. The first is that SWATH was not included in the comparison of Guo and Huan [2020a], whereas our study did include both AIF and SWATH. Secondly, while spectral matching to a library of known fragmentation spectra (e.g. GNPS, or MassBank) can be done, many compounds present in such databases have no matches thus reducing the identification rate observed from both DDA and DIA methods. Thus we introduce another dataset, constructed using an advanced multiple-injection method, to measure how well DDA and DIA methods perform with respect to exhaustive fragmentation procedures. Exhaustive fragmentation procedures have been shown to perform best with respect to coverage when a large number of replicates are available. However, real experiments are often constrained by cost or time, limiting the number of replicates that could be produced. In this setting, our analysis shows that DDA still performs best compared to DIA in recovering coverage when multiple replicates are not easily available.

Our recommendation on which acquisition method to choose is therefore:

- If the sample complexity is expected to be **low** to **medium**, OR it is preferred to **fragment as many features** as possible, even if they are not all identified, it is recommended to use DIA for data acquisition (keeping in mind the necessary deconvolution step).
- If the sample complexity is expected to be **high**, OR it is preferred to **obtain as many identified features** as possible, it is recommended to use DDA, which targets a specific ion for each MS2 scan thus generating fragmentation spectra that are almost immediately usable for analysis.

It is worth emphasizing that our results do not negate the usefulness of running DIA methods. Using DIA, we obtain the largest number of coverage of features, resulting in the most number of fragmented molecules – many of whom could be used for further investigation in the future. However acquired DIA scans need to be deconvoluted in order to translate the results to actual identification. Improvements in deconvolution methods are therefore needed to fully maximise the usefulness of DIA data. The work introduced here shows that a large margin of improvement is still possible in spectral deconvolution of DIA results – an avenue for further research that could be explored by the community. The work here also shows that through a simulation framework such as ViMMS, researchers could first test their deconvolution method in-silico. The portable nature of ViMMS means controllers (fragmentation methods) developed in simulators can be easily ported to run on the real instrument. DIA methods such as AIF and SWATH were implemented as controllers on top of ViMMS, tested in the simulator, and easily deployed to run on the actual instrument. This makes it easy for others to reproduce our results in simulation. It also opens the path for more advanced DIA methods to be developed in the future on top of ViMMS. In other fields such as molecular machine learning, having a standardized benchmark dataset [Wu et al., 2018] fosters development and collaborations as now a common reference exists onto which we can compare the performance of different methods. It is our hope that the simulated experiment introduced here could also serve as a standard and reproducible benchmark to which other deconvoluted methods can be compared to.

Our study is still limited in several aspects. The first is we only used MS-DIAL for the pre-processing of DIA data. MS-DIAL is de-facto the most popular tool used for spectra deconvolution and analysis of DIA data. However, other deconvolution tools exist, such as those of Yin et al. [2019] and Graca et al. [2022], that we do not include in our comparison. Parameters from MS-DIAL in our study were picked by hand to obtain reasonable results – similar to the limitation in Guo and Huan [2020a]. Furthermore, DDA and DIA methods exist that can exploit information across multiple samples [Koelmel et al., 2017, Tada et al., 2020] but these were not included in our evaluation. Using ViMMS, we could generate samples in multiple replicates and assess the performance of DDA and DIA methods when replicates are available. This validation to some extent has been done to compare standard DDA vs multiple samples/injections DDA in McBride et al. [2022] but not in the context of comparing to DIA methods.

For future work, the developed *in silico* benchmarking pipeline introduced in this work can also serve as the foundation to develop and validate a hybrid method that combines the benefit of both approaches. DDA and DIA methods exist in a spectrum: DDA isolates a certain precursor m/z for fragmentation, and DIA isolates multiple ions in a range of windows. This behaviour is the same across the entire run. It would be interesting to explore the potential of a hybrid method that combines the benefits of both DDA and DIA approaches. Such a method could potentially take advantage of the strengths of both approaches, allowing for more accurate and comprehensive analysis of complex samples. Such work has already been attempted [Guo et al., 2021] but that integration happens in data processing, not during the data acquisition itself. A hybrid acquisition method could use DDA to isolate certain prioritised precursor ions for fragmentation, and use DIA to isolate the remaining ions within a range of windows. This hybrid approach would allow for a more flexible and comprehensive analysis of samples, and could potentially improve the accuracy and reliability of results. This would lead to a more effective use of untargeted metabolomics, as more molecular features will be fragmented with higher-quality mass spectra associated to them. Ultimately, that will improve the biochemical interpretation of metabolomics profiles across the life sciences and other research areas.

## Author Contributions

JW led software development (in-silico and on the actual instrument), data acquisition and analysis. RMB and NT developed the evaluation codes for data analysis. SW performed sample preparation and instrument calibration. SR and VD conceptualised the research project. SR, RD and VD acquired funding. VD, KB and RD supervised the project. JJJvdH, KB, RD and SR performed manuscript review and editing. All authors contributed to the manuscript draft and the submitted version.

## Supplemental Data

Supplementary Materials are available as PDF. Python codes for the DDA and DIA controllers, as well as Jupyter notebooks to reproduce the results in this paper can be found in the ViMMS GitHub repository at https://github.com/glasgowcompbio/vimms, and under the following DOI: 10.5281/zenodo.7468914.

## Data Availability Statement

The dataset consisting of the DDA (Top-N, Intensity Non-overlap) and DIA (AIF, SWATH) mzML files generated and analysed for this study can be found in under the following DOI: 10.5525/gla.researchdata.1382.

