## Supplementary Information for "Simulated-to-real Benchmarking of Acquisition Methods in Metabolomics"

January 12, 2023

### 1 Supplementary Section S1 - Beer samples

The following a list of beer samples used:

| Index | Name | Type |
|---|---|---|
| 1 | Raspberry Sour by Vault City | Sour |
| 2 | Cacao & Hazelnut Broken Dream Twisted Breakfast Stout by Siren | Stout |
| 3 | Tennents | Lager |
| 4 | Life and Death by Vocation | IPA |
| 5 | Silence is... Citra by Overtone | Pale Ale |
| 6 | Punk AF by Brewdog | Alcohol-Free IPA |

Table 1: The list of beers used in the experiment.

#### 2 Supplementary Section S2 – MS-DIAL Parameters for Simulated Data

The following are MS-DIAL parameters used to process the DIA (SWATH, AIF) mzML files generated in simulation:

```
#Data type
MS1 data type: Centroid
MS2 data type: Centroid
Ion mode: Positive
DIA file: {config.txt}

#Data collection parameters
Retention time begin: 0
Retention time end: 10
Mass range begin: 0
Mass range end: 1100

#Centroid parameters
MS1 tolerance for centroid: 0.01
MS2 tolerance for centroid: 0.05

#Peak detection parameters
Smoothing method: LinearWeightedMovingAverage
Smoothing level: 3
Minimum peak width: 5
Minimum peak height: 1000
Mass slice width: 0.05

#Deconvolution parameters
Sigma window value: 0.5
Amplitude cut off: 10

#Adduct list
Adduct list: [M+H]+

#MSP file and MS/MS identification setting
MSP file: {msp_file}
Retention time tolerance for identification: 100
Accurate ms1 tolerance for identification: 0.025
Accurate ms2 tolerance for identification: 0.25
Identification score cut off: 0

#Text file and post identification (retention time and accurate mass based) setting
#Text file:
Retention time tolerance for post identification: 0.1
Accurate ms1 tolerance for post identification: 0.01
Post identification score cut off: 85

#Alignment parameters setting
```

```
Retention time tolerance for alignment: 0.05
MS1 tolerance for alignment: 0.0015
Retention time factor for alignment: 0.5
MS1 factor for alignment: 0.5
Peak count filter: 0
QC at least filter: True

#CorrDec setting
CorrDec excute: False
```

For AIF, {config.txt} is the following:

```
ID MS Type Start m/z End m/z Name CE DecTarget(1:Yes, 0:No)
0 SCAN 0 1100 0eV 0 0
1 ALL 0 1100 30eV 30 1
```

For SWATH, {config.txt} is the following:

```
Experiment MS Type Min m/z Max m/z
0 SCAN 0 1100
1 SWATH 0 100
2 SWATH 100 200
3 SWATH 200 300
4 SWATH 300 400
5 SWATH 400 500
6 SWATH 500 600
7 SWATH 600 700
8 SWATH 700 800
9 SWATH 800 900
10 SWATH 900 1000
11 SWATH 1000 1100
```

### 3 Supplementary Section S3 – MS-DIAL Parameters for Beer Samples

The following are MS-DIAL parameters used to process the DIA (SWATH, AIF) mzML files generated from actual beer samples produced on the Thermo instrument:

```
#Data type
MS1 data type: Centroid
MS2 data type: Centroid
Ion mode: Positive
DIA file: {config.txt}

#Data collection parameters
Retention time begin: 3
Retention time end: 24
Mass range begin: 70
Mass range end: 1100

#Centroid parameters
MS1 tolerance for centroid: 0.01
MS2 tolerance for centroid: 0.05

#Peak detection parameters
Smoothing method: LinearWeightedMovingAverage
Smoothing level: 3
Minimum peak width: 5
Minimum peak height: 25000
Mass slice width: 0.05

#Deconvolution parameters
Sigma window value: 0.5
Amplitude cut off: 10

#Adduct list
Adduct list: [M+H]+

#MSP file and MS/MS identification setting
#MSP file: {msp_file}
Retention time tolerance for identification: 100
Accurate ms1 tolerance for identification: 0.025
Accurate ms2 tolerance for identification: 0.25
Identification score cut off: 0

#Text file and post identification (retention time and accurate mass based) setting
#Text file: D:\Msdial-ConsoleApp-Demo files\Msdial-ConsoleApp-Demo files for DDA\Lipid_Nega_I
Retention time tolerance for post identification: 0.1
Accurate ms1 tolerance for post identification: 0.01
Post identification score cut off: 85

#Alignment parameters setting
```

```
Retention time tolerance for alignment: 0.16666666666
MS1 tolerance for alignment: 0.025
Retention time factor for alignment: 0.5
MS1 factor for alignment: 0.5
Peak count filter: 0
QC at least filter: True

#CorrDec setting
CorrDec excute: False
```

For AIF, {config.txt} is the following:

```
ID MS Type Start m/z End m/z Name CE DecTarget(1:Yes, 0:No)
0 SCAN 70.0 1100.0 0eV 0 0
1 ALL 70.0 1100.0 25eV 25 1
```

For SWATH, {config.txt} is the following:

```
Experiment MS Type Min m/z Max m/z
0 SCAN 70.0 1100.0
1 SWATH 70 170
2 SWATH 170 270
3 SWATH 270 370
4 SWATH 370 470
5 SWATH 470 570
6 SWATH 570 670
7 SWATH 670 770
8 SWATH 770 870
9 SWATH 870 970
10 SWATH 970 1070
```

### 4 Supplementary Section S4 – Intensity Non-overlap Acquisition Method

*Intensity Non-overlap* is an advanced iterative DDA-based method that incorporates several new concepts to achieve more targeted fragmentations and prevent more redundant ones across samples or injections. This incorporates several key ideas to greatly increase the fragmentation coverage of unique molecular features across multiple injections. First is the method incorporates the concept of tracking Region of Interest (RoI, groups of related precursor ions) in real-time and prioritising RoIs for fragmentation rather than individual ions. Then across repeated injections, the same RoIs if seen again are downweighted by their non-overlapping area. Finally its score component also prioritises acquiring the same RoI again if it can be fragmented at a higher apex than before as compared to the previous injection. Intensity Non-overlap is builds upon 'TopNEXt' [1], a real-time scan prioritisation framework and an extension of the Virtual Metabolomics Mass Spectrometer which implements several improved multi-sample fragmentation strategies within a modular and cohesive base. For more details, please refer to [1].

#### References

[1] Ross McBride, Joe Wandy, Stefan Weidt, Simon Rogers, Vinny Davies, Rónán Daly, and Kevin Bryson. TopNEXt: Automatic DDA exclusion framework for multi-sample mass spectrometry experiments. *In Submission*, 2022.
